# Extreme resistance to *Potato Virus Y* in potato carrying the *Ry_sto_* gene is mediated by a TIR-NLR immune receptor

**DOI:** 10.1101/445031

**Authors:** Marta Grech-Baran, Kamil Witek, Katarzyna Szajko, Agnieszka I Witek, Karolina Morgiewicz, Iwona Wasilewicz-Flis, Henryka Jakuczun, Waldemar Marczewski, Jonathan DG Jones, Jacek Hennig

**Affiliations:** Institute of Biochemistry and Biophysics, Polish Academy of Sciences, Pawińskiego 5a, 02-106 Warsaw, Poland; The Sainsbury Laboratory, University of East Anglia, Norwich Research Park, Norwich, NR4 7UK, UK; Plant Breeding and Acclimatisation Institute-National Research Institute, Platanowa 19, 05-831 Młochów, Poland

## Abstract

*Potato virus Y* (PVY) is a major potato pathogen that causes annual losses of billions of dollars. Control of its transmission requires extensive use of environmentally damaging insecticides. *Ry_sto_* confers extreme resistance (ER) to PVY and is a valuable trait in resistance breeding programs. We isolated *Ry_sto_* using Resistance gene enrichment sequencing (RenSeq) and PacBio SMRT (Pacific Biosciences Single-Molecule Real Time Sequencing). *Ry_sto_* encodes a nucleotide binding-leucine rich repeat (NLR) protein with an N-terminal TIR domain, and is sufficient for PVY perception and extreme resistance in transgenic potato plants. We investigated the requirements for *Ry_sto_*-dependent extreme resistance, and showed that *Ry_sto_* function is temperature-independent and requires EDS1 and NRG1 proteins. Ry_sto_ may prove valuable for creating PVY-resistant cultivars of potato and other *Solanaceae* crops.

## INTODUCTION

*Potato virus Y* (PVY), a member of the *Potyvirus* genus (Potyviridae family), is the most economically important virus infecting potato crops, with tuber yield and quality losses ranging from 10 to 90 % depending on year, cultivar and location (De Bokx and van der Want 1987; Valkonen 2007). PVY can also infect other *Solanaceae* species such as bell pepper, tomato and tobacco (Aramburu 2006). The virus is easily transmitted mechanically by over 40 aphid species in a non-persistent manner (Brunt 2001). Intracellular pathogens such as viruses are difficult to control chemically; PVY outbreaks are addressed by the destruction of infected plants or pesticide treatment to limit the population of vectors (Kopp et al. 2015). Resistance breeding remains the best method to prevent viral infections. PVY variants are highly variable at the biological, serological, and molecular levels (Scholthof *et al.* 2011). They can be classified into different strain groups depending on host response. Virus strains PVY^0^ and PVY^C^ usually induce mild symptoms of infection, e.g. leaf mosaic lesions, crinkling, leaf drop and dwarfing, while leaf symptoms of PVY^N^ and PVY^N^W infection are barely noticeable. In addition to severe leaf symptoms, PVY^NTN^ infection leads to potato tuber necrotic ringspot disease which greatly affects tuber marketability (Schubert et al. 2007).

Plant defense responses to viruses are multifaceted and depend on several factors including plant age and tissue type. The most durable control of PVY is provided by resistance genes. In potato, there are two main types of resistance against PVY: extreme resistance (ER), conferred by the *Ry* genes, and hypersensitive resistance (HR), conferred by the *Ny* genes (Valkonen 2015). The HR to PVY is usually strain-specific and may result in a range of necrotic reactions in both locally and systemically infected leaves (Valkonen et al. 1998). However, under some conditions HR may be ineffective for restriction of PVY in plants (Vidal et al. 2002). In contrast ER genes are broad-spectrum and confer strong and durable resistance, characterized by lack of visible symptoms after inoculation (Flis et al. 2005).

Resistance genes against PVY infection were introduced into potato cultivars from wild or domesticated *Solanum* species. Ten genes for resistance to PVY and one gene for resistance to PVA (a potyvirus related to PVY) were mapped to four potato genome segments on chromosomes IV, IX, XI and XII (Valkonen et al., 2017, van Eck et al. 2017). Some of these genes, for instance *Ry_sto_* and *Ry-f_sto_* from *S. stoloniferum*, were introduced into multiple potato cultivars worldwide and shown to confer durable resistance against multiple PVY strains. Nevertheless, no genes conferring effective HR or ER-type of resistance have previously been isolated (Valkonen et al. 2017).

It has been estimated that controlling major diseases through the informed deployment of resistance genes could contribute over 30% towards crop yield whilst reducing the requirements for chemical applications (Gebhardt and Valkonen 2001). This might be achieved by implementation of new breeding tools to select resistant genotypes and deploy them as varieties (Armstrong et al 2018).

Most plant resistance genes encode intracellular nucleotide-binding, leucine-rich repeat (NLR) receptors, some of which carry an N-terminal TIR domain (TIR-NLRs), and others an N-terminal coiled coil domain (CC-NLRs) (Jones et al., 2016). A useful tool for identification and assembly of full length NLR-encoding genes that cosegregate with resistance is SMRT RenSeq (Witek et al. 2016). The methodology is more robust and cost-effective in monitoring NLR sequences than whole-genome sequencing. All currently known NLRs effective against viruses, nematodes and the late blight pathogen *Phytophthora infestans* can be tracked with SMRT RenSeq in potato and their polymorphisms defined (Armstrong et al 2018).

There is a high demand for strong and durable genes against PVY infection and *R* genes conferring ER to PVY fulfil this requirement. As the understanding of mechanisms that underlie ER response to viruses is still incomplete it is of key importance to identify the network of factors that contribute to effective resistance against the pathogen.

In this study, we set out to isolate the *Ry_sto_* gene conditioning the ER trait using SMRT RenSeq. We found multiple TIR-NLR-encoding paralogs that cosegregate with resistance, and demonstrated that paralog *Ry_sto_* confers resistance to PVY and to the related potyvirus PVA in different *Solanaceae* backgrounds. We show that PVY and PVA coat proteins are recognized as *avr* factors for *Ry_sto_* triggered immunity. Finally, we investigate the functional relationship between ER and HR, and show that extreme resistance is EDS1, NRG1 but not SA or temperature dependent.

## RESULTS

### *Ry_sto_* originates from *S. stoloniferum*

Extreme resistance to PVY was introgressed to commercial potato cultivars from three different wild *Solanum spp*. In the European cultivars, resistance originates mostly from *S. stoloniferum* (*Ry_sto_*) and is not yet overcome (Valkonen et al. 2017). Therefore, we set out to clone *Ry_sto_* using dihaploid clone (dH) Alicja, which has PVY resistant *S. stoloniferum* in its ancestry. We generated diploid mapping population consisting of 391 F1 individuals, and evaluated for resistance to PVY by mechanical and graft inoculations as described in Flis *et al*. (2005). The segregation ratio of resistant versus susceptible progeny in the mapping population deviated from the 1:1 ratio expected for the segregation of a single dominant gene and was distorted towards resistance (149 susceptible and 242 PVY-resistant F1 individuals).

### RenSeq combined with SMRT sequencing and bulked segregant analysis leads to successful cloning of *Ry_sto_*

It was previously shown that *Ry_sto_* maps to the distal end of long arm of potato chromosome XII (Flis *et al* 2005, Song *et al* 2005, van Eck *et al* 2017), which corresponds to the region downstream of 58 Mb in the reference potato genome clone DM (PGSC, 2011). Jupe *et al.* (2013) showed that this region contains 18 complete and partial NLR immune receptors from both TIR-NLR and CC-NLR classes, organized in four clusters (Figure 1a). We therefore hypothesized that *Ry_sto_* likely encodes an NLR, and applied RenSeq to predict candidate genes. To assemble the NLR-encoding genes of the resistant parent with high confidence, SMRT-RenSeq was performed. Assembly of PacBio Reads of Interest (ROI) resulted in 1,555 contigs, out of which 1254 were annotated as NLRs using NLR-parser software (Steuernagel *et al.* 2016). Mapping of short read RenSeq data from resistant (R), susceptible (S) and bulked susceptible (BS) plants, and subsequent polymorphism calling resulted in 10 linked contigs showing presence/absence polymorphism. Expression of candidate genes was confirmed using cDNA-RenSeq data obtained from the resistant parent. These analyses resulted in 11 candidate NLRs. Four candidate genes belonged to CC-NLR and seven to the TIR-NLR class. A phylogenetic tree constructed with full-length amino acid sequences showed that candidate genes belong to three different clades and show only distant relation with cloned functional *Solanaceae* genes. All TIR-NLRs belonged to the same clade and show amino acid identity of between 75% and 90%. The CC-NLRs genes split into two distinct clades, consistent with their physical separation in the potato genome (Fig. 1, Supplementary Fig. 1).

**Figure 1:**
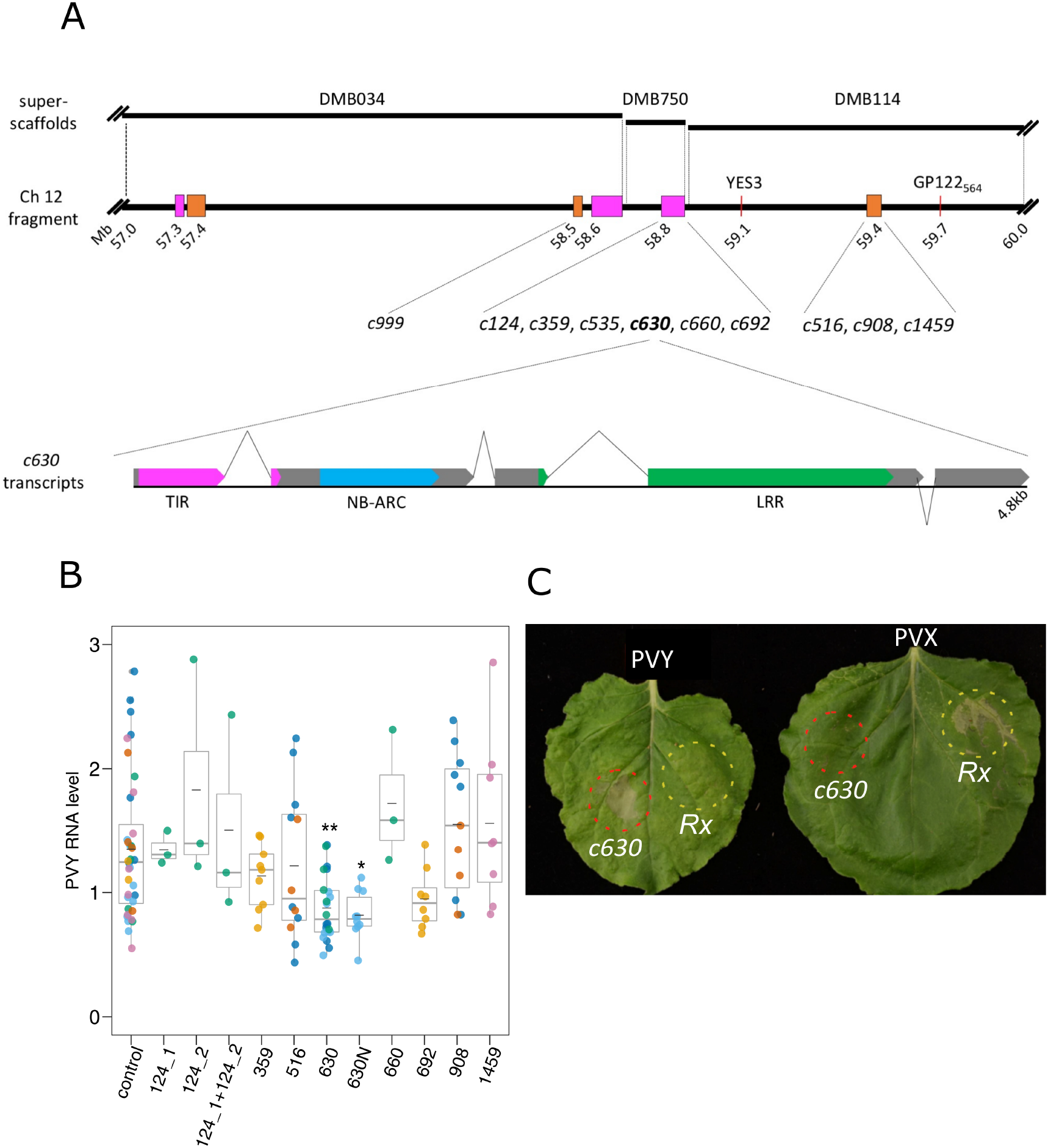
Functional analysis of candidate genes in transient expression assay in *N. benthamiana* plants. (a) Schematic representation of candidate genes and a fragment of chromosome 12 linked to the *Ry_sto_* gene. Potato superscaffolds (DMB034, DMB750 and DMB114; top panel) were aligned to the distal end of longer arm of potato chromosome 12 (57 to 60Mb fragment shown; middle panel), which is linked to the *Ry_sto_* gene. Annotated NLRs/NLR clusters are depicted in orange (CNL) and magenta (TNL). Red lines indicate known markers linked to *Ry_sto_*. Distances are given in megabases (Mb) according to the DM potato reference genome (PGSC 2011), numbers indicate proximal position of marked sequences. Candidate contigs for *Ry_sto_* derived from SMRT RenSeq were aligned to the DM potato genome and best hit position is shown on the map. Map drawn to the scale. Lower panel presents schematic structure of the *c630* transcripts. Most abundant version (supported by ~80% of cDNA-RenSeq reads) consists of 4 exons. Additional intron (depicted with inverted lines) positioned 24 nucleotides upstream of initial stop codon results in fifth exon at 3’ end. Exons drawn in grey, colors indicate canonical NLR domains; TIR (magenta), NB-ARC (blue) and LRR (green). Scheme drawn to the scale. (b) Relative level of PVY RNA after infection of *N. benthamiana* leaves transiently expressing candidate contigs. Three leaves of *N. benthamiana* plants were infiltrated with the vector pICSLUS0003::35S overexpressing *c124_1*, *c124_2*, *c359*, *c516*, *c630*, *c660*, *c692*, *c908*, and *c1459*, pICSLUS0001 vector overexpressing c630 under control of native regulatory elements or an empty vector48 hours later leaves were inoculated with PVY^NTN^ strain. Seven days after PVY inoculation mRNA was isolated from upper, non-inoculated leaves and PVY RNA levels were quantified with qPCR. As reference genes *EF1* and *L23* were used. Error bars represent the standard deviation of the means (-). Additionally median is also presented. Statistical analysis was perform using one-way ANOVA with Dunnett’s test. Experiments were performed on 3-10 plants for each construct and repeated three times. (c) HR response of *N. benthamiana* plants expressing *c630* or *Rx* genes. Two fully developed leaves of *N. benthamiana* plants 14 days following PVY^NTN^ (left) or PVX infection (right), were infiltrated with a suspension of *A. tumefaciens* carrying pICSLUS0003*c630* (area marked with red) or pGBT-Rx-GFP (area marked with yellow) Seventy-two hours after infiltration expression of *c630* in PVY infected plants resulted in HR similarly to when *Rx* gene was delivered into plants infected with PVX virus. Experiment was repeated three times.

Homology searches of candidate genes to the reference genome positioned them in three clusters located in the distal end of the long arm of chromosome 12 (58-59Mb of DM reference genome, Fig. 1a), where markers linked to *Ry_sto_* are also located (GP122_564_, Witek *et al*. 2006 and YES3, Nie *et al.* 2016). This part of chromosome 12 consists of three neighboring superscaffolds DMB034, DMB750 and DMB114 and one of them (DMB114) was suggested to be a likely location of *Ry_sto_* (van Eck *et al*. 2017). Thus, all selected candidates NLR genes from SMRT RenSeq data localize to the distal end of chromosome 12 where *Ry_sto_* was previously positioned.

### Functional analysis of candidate contigs in transient expression assay in *N. Benthamiana* plants

To establish whether the candidate genes are functional *in planta*, the open reading frames (ORFs) of 11 co-segregating NLRs were PCR-amplified and placed under control of the *35S* promoter in a binary vector pICSLUS0003 as described (Witek et al., 2016). Two functional tests were then performed. In the first test, the constructs were transiently expressed in *N. benthamiana*, followed by PVY^NTN^ inoculation. At 7 dpi, leaf samples were collected and levels of viral RNA were measured using qPCR. One candidate gene (*c630*) reduced virus multiplication as compared to control *N. benthamiana* plants (Fig. 1b). The *c630* ORF with its native 5’ and 3’ regulatory sequences (2,263 nt upstream from 5’ and 1,013 nt downstream from 3’ end, respectively) was cloned into the pICSLUS0001 binary vector as described (Witek et al., 2016). Transient assays in *N. benthamiana* also showed a statistically significant (*P* = 0.011475) inhibition of virus multiplication (Fig. 1b). Candidate gene *c999* showed autoactivity, even when infiltrated with low OD (0.1) and another candidate gene, *c535* could not be cloned.

In the next step, *N. benthamiana* plants were systemically infected with PVY^NTN^ or PVX, followed by *c630* transient expression by Agro-infiltration. The expression of *c630* in plants carrying PVY^NTN^ resulted in a hypersensitive response (HR) (Fig. 1c), similar to when the *Rx* gene was delivered into plants infected with PVX (Bendahmane et al., 1999)

### Expression of *c630* triggers HR to different viruses

To test whether *c630* elicited an immune response to different of strains of PVY *N. benthamiana* plants were infected with: 0, N, N-Wilga, NTN isolates of PVY, or cross related Potato Virus A (PVA). At 7 dpi, leaves showing symptoms of viral infection were infiltrated with *Agrobacterium* strains carrying *c630* or an empty vector. Transient expression of *c630* resulted in a strong HR in plants carrying all PVY isolates and PVA, but not PVX (Potato virus X) or TMV (Tobacco Mosaic Virus) (Supplementary Tab. 1.). No HR was observed when *c630* was not expressed in control plants. These experiments further verified *c630* as a candidate for the functional *Ry_sto_* gene.

### *Ry_sto_* expression restricts systemic spread of PVY and PVA

To further investigate *Ry_sto_* function, stable transgenic *Solanaceae* plants (potato and tobacco) carrying *Ry_sto_* or two other non-functional homologues under the control of *35S* promoter or native regulatory elements were created using *Agrobacterium-*mediated transformation.

Two potato cultivars *S. tuberosum* cv. Maris Piper (MP) and cv. Russet Burbank (RB), susceptible to PVY, were then chosen to test *Ry_sto_* functionality. Transgenic plants from both cultivars carrying *Ry_sto_* under control of *35S* promoter or native promoter (for MP only) were inoculated with PVY^NTN^ or mock treated with water. Systemic virus spread was monitored in upper, non-inoculated leaves using enzyme-linked immunosorbent assay (ELISA) at 3-weeks after inoculation. In the MP background expressing Ry_sto_ under control of *35S*, in 10 out of 12 tested transgenic lines PVY was not detected in the upper part of the plant (Supplementary Tab. 2). To confirm these ELISA results, three randomly chosen transgenic lines (two resistant and one susceptible) were tested using qPCR (Fig. 2b). The inhibition of virus multiplication and spread in two resistant transgenic plants correlated with elevated levels of *Ry_sto_* gene expression. In the susceptible transgenic line, the expression of *Ry_sto_* transgene was not detected. In cv. Russet Burbank in all 4 tested independently isolated transgenic lines, PVY was not detected using ELISA or using qPCR (Fig. 2c, Supplementary Tab. 3).

**Figure 2:**
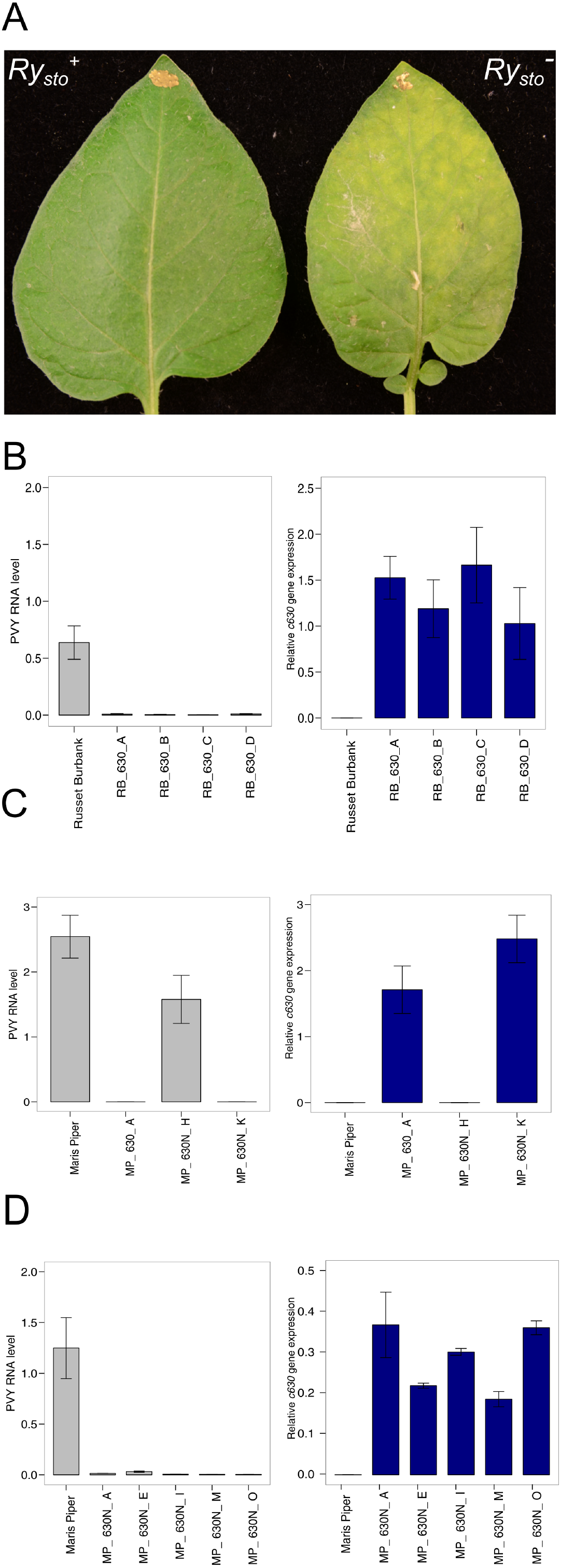
Expression of *Ry_sto_* leads to immunity of transgenic plants to PVY infection. (a) Illustration of ER type response of PVY-inoculated leaves of *S. tuberosum* cv Russet Burbank plants expressing *Ry_sto_.* Four-week-old transgenic potato plants cv. Russet Burbank carrying construct *35S::Ry_sto-630_*, and non-transformed plants were inoculated with PVY^NTN^. Two weeks after infection chlorosis were observed on inoculated leaves of non-transformed plants (right) in contrast *Ry_sto_* transgenic plants (left) remained symptomless. (b) Effect of *Ry_sto_* expression completely blocks PVY spreading in transgenic potato plants. Four-week-old transgenic potato plants cv. Russet Burbank (panel B) or Maris Piper (panel C) carrying construct *35S:: Ry_sto_*, Maris Piper plants carrying *Ry_sto_* construct under control of native 5’- and 3’-regulatory elements (panel D) and non-transformed plants were inoculated with PVY^NTN^. Three weeks after viral inoculation, mRNA was isolated from upper, non-inoculated leaves. The PVY RNA levels and the expression of *Ry_sto_* were quantified using qPCR, relative to the *EF1* and *Sec3* reference genes and expressed as means ± SD calculated from three biological replicates per plant line.

Additionally, when *Ry_sto_* gene was expressed under control of its native promoter all tested lines (5 lines) were PVY free (Fig. 2d). In all resistant lines non-macroscopic symptoms of HR were observed, regardless of the genotype of potato used (Fig. 2a).

*N. tabacum 35S::Ry_sto_* transgenic and non-transformed control plants were inoculated with PVY^NTN^ as previously. At 5 dpi, some necrotic lesions were visible only on inoculated leaves of *Ry_sto_* plants, while no macroscopic symptoms of HR were observed on the non-transformed control plants (Supplementary Fig. 2). In addition, control susceptible plants developed typical systemic symptoms of PVY infection. qPCR on upper, non-inoculated leaves showed that systemic spread of the virus was fully restricted in *Ry_sto_* expressing lines, while PVY RNA levels in non-transformed plants were high. These results were confirmed in T0 (Supplementary Fig. 3A) and T1 generation plants (Supplementary Fig. 3B). In transgenic tobacco lines expressing *Ry_sto_* under control of its natural 5’ and 3’ regulatory elements, a local HR similar to that seen in *35S::Ry_sto_* plants was also observed at 5 dpi. The absence of viral RNA was confirmed with qPCR (Supplementary Fig. 4) at 7 and 14 dpi and was fully correlated with *Ry_sto_* transcript expression.

Introduction of *Ry* genes from *Solanum stoloniferum* to *S. tuberosum* may condition resistance not only to PVY but also to close related PVA (Cockerham, 1970). To test whether expression of *Ry_sto_* determines resistance to PVA, tobacco plants with and without functional *Ry_sto_* transgene were inoculated with this virus. Western blotting analysis with anti PVA antibodies revealed inhibition of PVA systemic spread in *Ry_sto_* transgenic ones (Supplementary Fig. 5).

### The PVY and PVA coat proteins (CP) are elicitors of Ry_sto_ triggered immunity

It was previously reported that there is a single ORF encoding a polyprotein in the PVY genome, which is cleaved by three viral proteases to form functional proteins (Jakab et al., 1997). Thus, to identify the elicitor of the *Ry_sto_*-triggered immunity the open reading frame from the most promising putative viral proteins (HcPro, NIa, NIb, coat protein [CP]) were cloned and transiently expressed with in transgenic *Ry_sto_* and non-transformed tobacco plants. Among all tested proteins only the coat protein induced strong HR at 2-3 days post treatment in *Ry_sto_* plants, whereas no HR was observed in WT control (Fig. 3a). A similar type of analysis was performed for PVA virus. Only PVA coat protein was able to elicit HR when is expressed in transgenic *Ry_sto_* plants (Fig. 3b).

**Figure 3:**
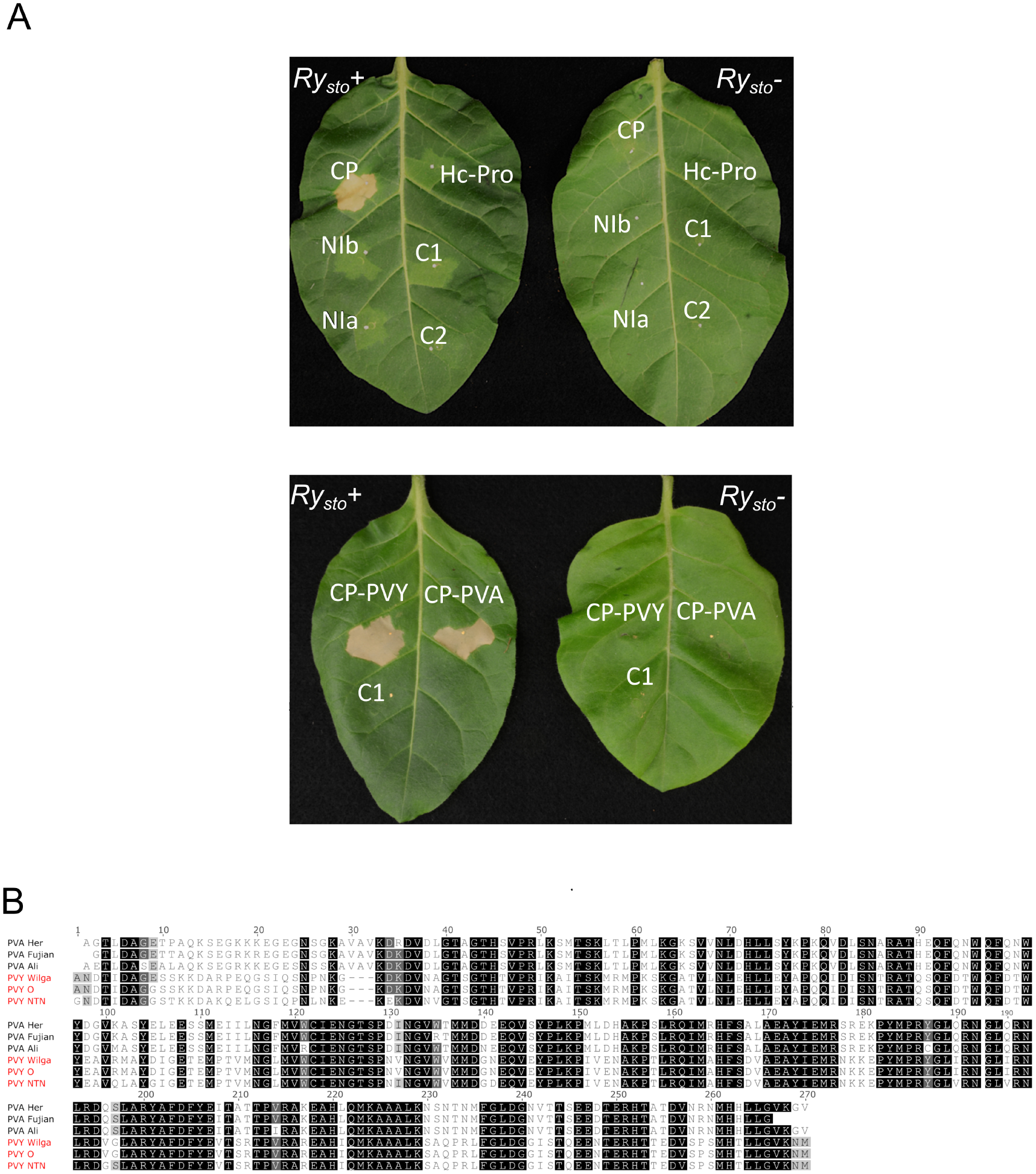
Coat proteins of two close related potyviruses elicit of *Ry_sto-_*triggered immunity. (a) Ry_sto_ recognizes (upper panel) PVY and (lower panel) PVA coat proteins as avirulence factors in transient expression assays. To identify the elicitor of the *Ry_sto_* mediated resistance against PVY or PVA viruses the open reading frame of putative viral proteins (HC-Pro, Nia, Nib and CP) was cloned to pBIN53 vector and transiently expressed in *N. tabacum. Ry_sto_* transgenic and non-transformed plants. As controls (C1, C2) *A. tumefaciens* or infiltration buffer were used. Three days after treatment HR was observed only for PVY or PVA coat protein (CP). (b) Comparison of the selected PVY and PVA coat protein amino acid sequences. The comparison is made against PVY^NTN^ CP. Identical residues are shaded in black. The PVA CP shares above 59% identity with the PVY sequence. The alignment was generated using MAFFT-L-INS-I (Katoh and Toh, 2008) and visualized in Jalview 2,10,4b1 (Waterhouse et al. 2009).

### Extreme resistance confirmed by *Ry_sto_* is epistatic to the Ny-1 mediated HR

The functional relationship between two types of defence response to PVY infection, ER and HR, was investigated. Potato cultivar (Rywal) carrying the *Ny-1* gene, responsible for HR to PVY infection (Szajko et al., 2008 and Baebler et al., 2014) was transformed with *Ry_sto_*. Plants of both genotypes (*Ny-1* or *Ry_sto_* /*Ny-1*) were inoculated with PVY^NTN^ as described previously. Five days after infection HR developed only in parental genotype (*Ny-1*), while no sign of HR was observed in the *Ry_sto_* /*Ny-1* genotype (Fig. 4b). This indicates that ER conferred by *Ry_sto_* is epistatic to *Ny-1* dependent HR.

**Figure 4:**
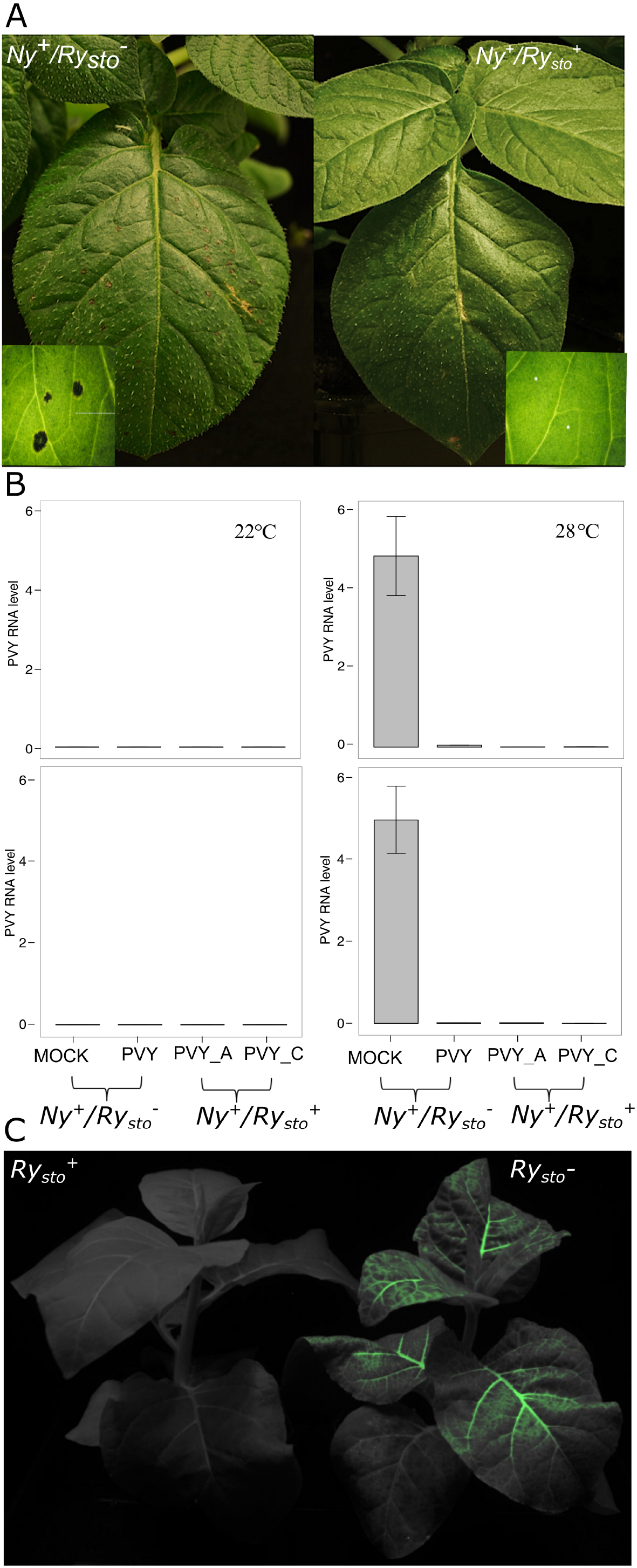
*Ry_sto_* dependent ER is epistatic to the HR and temperature independent. (a) Determination of epistatic effect of ER to HR. Potato *Ry_sto_*/*Ny-1* and *Ny-1* plants were inoculated with PVY^NTN^. Five days after infection symptoms of local cell death occurred only in *Ny-1* plants. Experiment was repeated three times. (b) *Ry_sto_* dependent PVY multiplication is inhibited in elevated temperature. Potato *Ry_sto_* /*Ny-1* and Ny-1 plants were inoculated with PVY^NTN^ and divided to two groups; the first group was kept at 20°C, the second at 28°C. Water-treated plants were used as negative controls. Seven days following inoculation, samples of inoculated (lower graphs) and upper non-inoculated leaves (upper graphs) were collected and PVY RNA levels were measured qPCR, relative to the *EF1* and *Sec3* reference genes and expressed as means ± SD calculated from three biological replicates per plant line. A and C describes names of transgenic *Ry_sto_* lines used. (c) Stable transgenic *N. tabacum* plants carrying *Ry_sto_* under control of a 35S promoter are resistant to PVY infection in elevated (32°C) temperature. Seven-week-old *N. tabacum 35S::Ry_sto_* transgenic and non-transformed plants were inoculated with PVY^N205-^GFP clone. Typical symptoms of PVY infection were observed after 7 dpi in non-transformed plants in contrast *Ry_sto_* lines remained symptomless. Picture was taken at 14 dpi.

### Extreme resistance mediated by is temperature-independent

To investigate the role of temperature in *Ry_sto_*-mediated ER, the response to PVY infection was studied in a *Ny-1* or *Ry_sto_* /*Ny-1* genotype. PVY mRNA levels were measured 14 dpi in upper non-inoculated leaves following mock and virus treatments in two types of temperature conditions (22°C and 28°C). In both genotypes, no PVY multiplication was observed when plants were kept at 22°C. In contrast, when kept in elevated temperature in *Ny-1* genotype HR was inhibited and PVY RNA was detected systemically, whereas in transgenic *Ry_sto_* /*Ny-1* lines PVY spreading was still fully inhibited (Fig. 4a,b). To test whether *Ry_sto_* hinders systemic infection above 28°C, transgenic and non-transformed tobacco plants were infected with GFP-tagged PVY (PVY^N605^-GFP) at 32°C. Interestingly, expression of *Ry_sto_* at higher temperature still prevented systemic spread of the virus (Fig. 4c).

These observations suggest that unlike the tobacco N gene, which also encodes a TIR-NLR, high (>28°C) temperature does not compromise *Ry_sto_* immunity to PVY infection (Samuel 1931).

### NRG1 and EDS1 but not SA are crucial for Ry_sto_ dependent immunity

For N-dependent resistance of tobacco to TMV, additional genes EDS1 and NRG1 are required (Collier *et al.*2011, Liu et al., 2002, Peart et al., 2005). Like N, *Ry_sto_* carries a TIR domain, so we examined whether these two regulator proteins mediate a Ry_sto_ dependent response to PVY infection. To test this hypothesis, *EDS1* or *NRG1 N. benthamiana* mutant plants were infected with PVY^NTN^ followed by transient expression of *Ry_sto_* or *GFP*, as a control. 72 hpi in WT plants, HR was observed. In contrast, both knockout plants remained symptomless. This suggests that both NRG1 and EDS1 proteins are crucial for the Ry_sto_-mediated response. A similar analysis was performed with *N. benthamiana* plants expressing bacterial *NahG* that encodes salicylate hydroxylase (Friedrich et al., 1995). In WT and in *NahG* plants, symptoms of HR were observed at 72 hpi (Fig. 5). This suggests that elevated levels of SA are not essential for establishing a Ry_sto_ dependent resistance.

**Figure 5:**
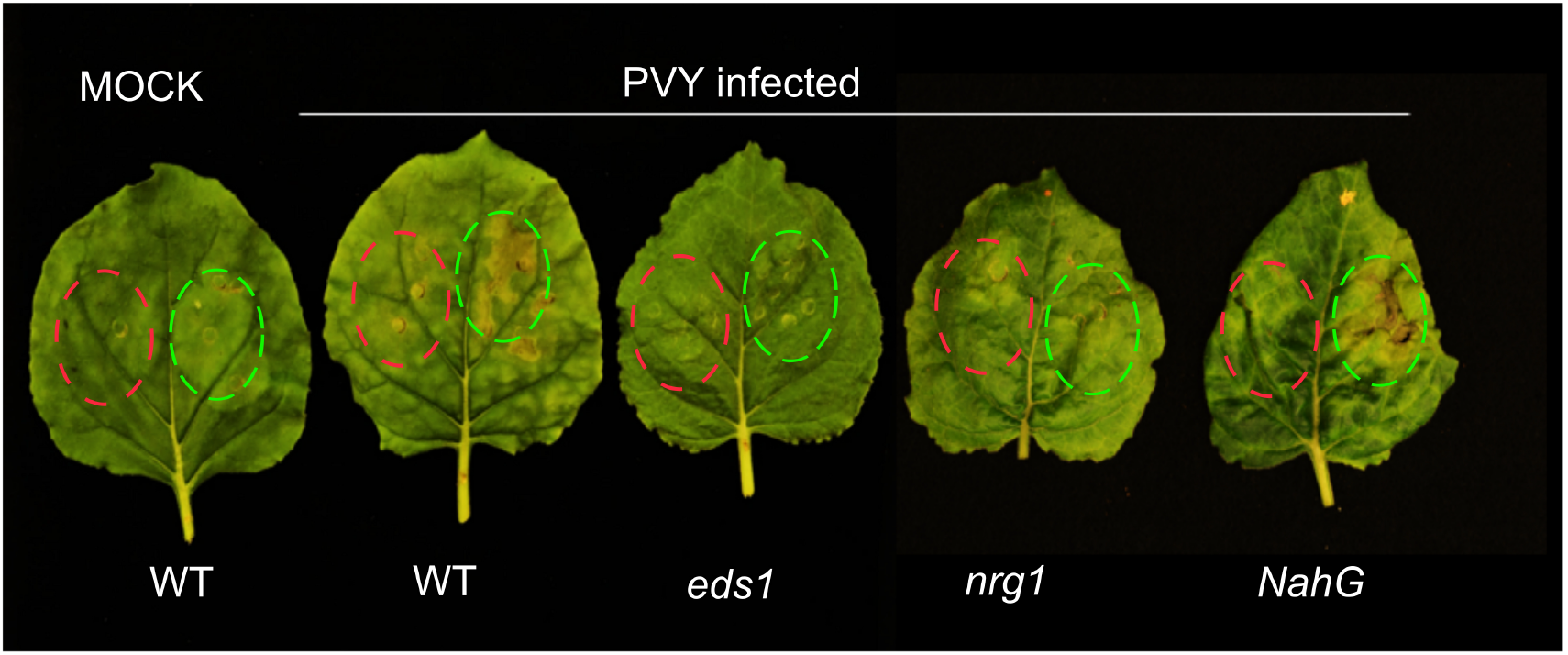
Downstream signaling components crucial for *Ry_sto_-*mediated immunity. Fully developed leaves of *N. benthamiana EDS1, NRG1* knockout plants or *NahG* and non-transformed ones were infected with PVY^NTN^ or Mock-treated. Two weeks later leaves with symptoms of PYV infection were infiltrated with suspension of *A. tumefaciens* carrying pICSLUS0001:*c630* (highlighted in green) or pGBT-*GFP* (highlighted in red) as a control. Three days after infiltration in non-transformed plants and *NahG* plants (SA free) symptoms of cell death were observed. The experiment was repeated three times with similar results.

## DISCUSSION

Genes conferring ER against PVY infection had been introduced into potato cultivars from wild or domesticated species from *Solanaceae* family, e.g. *S. chacoense, S. stoloniferum* or *S. tuberosum* ssp*. andigena*. Two alleles of *Ry_sto_* gene derived from *S. stoloniferum*: *Ry*-*f_sto_* and *Ry_sto_*, have most often been used (Flis et al. 2005, and Song et al. 2005). In this study, we report the isolation and characterization of a novel, broad-spectrum resistance gene *Ry_sto_* using RenSeq (Jupe et al. 2013, Witek et al. 2016).

We hypothesized that the underlying PVY resistance gene encodes an NLR protein. Data obtained from SMRT and Illumina sequencing together with SNP calling and presence/absence polymorphism detection revealed 11 transcriptionally active NLR genes, as candidates for the *Ry_sto_* gene. We mapped these candidates to the DM reference genome (PGSC 2011) at the distal end of chromosome 12, a region shown to carry *Ry_sto_*-linked markers. Additionally, using graphical genotyping on panel of tetraploid potatoes, van Eck et al. (2017) suggested that *Ry_sto_* locates on super-scaffold DMB114. This is consistent with our data, as three of 11 candidates were located on this super-scaffold, while remaining on neighboring super-scaffolds, all within 1 Mb distance. Interestingly, all linked NLRs showed presence/absence polymorphism with the same allele ratio, indicating that the whole interval introgressed from *S. stoloniferum* rarely recombines in cultivated potato.

Functional assays in *N. benthamiana* and cultivated potato plants confirmed that only one candidate, *Ry_sto_*, stopped PVY multiplication and spread. The protein encoded by *Ry_sto_* has motifs and domains typical of TIR-NLR-type resistance proteins and shares less than 34% identity at amino acid level with previously described TNL-type resistance proteins from *Solanaceae*., including: *N*, *Y-1*, *Bs4* and *Pvr4*.

As most potato cultivars lack broad-spectrum resistance to circulating strains of PVY (Valkonen 2015) we showed that *Ry_sto_* confers resistance against multiple PVY strains (Suppl. Table 1). In contrast, for *Rx*, *Rsv3 or Sw*-*5* genes mediating ER to other viruses, isolates breaking immunity have appeared (Moreira et al. 1980, Querci et al. 1995, Brommonschenkel et al. 2000, Zhang et al. 2012). Recently two novel genes conferring immunity to some PVY strains were described. Kim et al (2017) isolated, from *Capsicum annuum*, a dominant gene *Pvr4* conferring a broad range of resistance to potyviruses, including some PVY isolates. However, it was tested only in heterologous system (*N. benthamiana*) and not in any *Solanaceae* crops. A single recessive gene, called *NtTPN1*, isolated from tobacco causes veinal HR-like necrosis (Michel et al., 2018). Moreover, *Pvr4* was never tested in potato and there is no evidence that it would be functional in that background. Our results suggest that the *Ry_sto_*-mediated immunity is effective in several potato cultivars and tobacco not only against different strains of PVY but also to the related PVA (Fig. 2a, 4c, Supplementary Fig. 4).

We tested multiple PVY reading frames and found that viral coat protein (CP) is the recognized factor in Ry_sto_-mediated immunity against PVY and PVA (Fig. 3a). PVY and PVA CPs share ~59% identity (Fig. 3b) and have nucleocytoplasmic localization in infected cells (Supplementary Fig. 6). Interaction of Ry_sto_ with CP, directly or indirectly, initiates a response leading to ER by preventing virus accumulation at an early stage of infection. Resistance may be due to inhibition of viral cell-to-cell movement *via* callose deposition as was reported in soybean-SMV pathosystem (Seo et al., 2014) or by translational arrest of viral transcripts as was described for *N* gene-mediated resistance to tobacco mosaic virus (TMV) in a process that involves Argonaute 4 (Bhattacharjee et al., 2009). The CP might be recognized Ry_sto_ either by recognition of specific amino acid sequence or of a conserved element of CP structure as was reported for Rx protein (Baurès et al., 2000, Candresse et al., 2010).

Rx can trigger HR when co-expressed with PVX coat protein (Bendahmane et al. 1999). Using *N. benthamiana* we found that *Ry_sto_* behaves similarly. When plants were infected with PVY followed by *Ry_sto_* transient expression a strong HR was observed (Fig. 1c). We speculated that the nature of the *Ry_sto_* response, as either ER or HR, might be determined by the elicitor expression levels or genetic background used. When CP was expressed under control of constitutive promoter (35S) in transgenic *Ry_sto_* tobacco plants it elicited an HR, whereas CP produced during PVY infection led to ER. Moreover, when *Ry_sto_* was introduced into *S. tuberosum* plants no HR symptoms were observed but in *N. tabacum*, necrotic lesions on inoculated leaves were noticed upon PVY infection (Supplementary Fig. 2). This was also observed for *Rx* gene where the specificity of recognition was modulated by host factors differing between *Rx*-expressing potato and tobacco plants (Baurès et al., 2008).

For different resistance genes, ER or HR formation and control of viral spread might be determined by the dose of the R gene. High level *HRT* gene expression in susceptible *Arabidopsis thaliana* leads to restriction of virus replication or movement without any visible symptoms of HR, while lower levels of HRT transcript in resistant *Arabidopsis* lines leads to micro-HR or HR with systemic virus movement, respectively (Cooley et al., 2000). In contrast, our results suggest that ER is not affected by *Ry_sto_* expression levels. Relative expression levels of *Ry_sto_* differ between lines depending on the promoter used. Transgenic potato lines under control of native regulatory elements exhibited much lower transcript expression in comparison to those under control of 35S. Nevertheless, these differences did not influence the ER phenotype which was observed in all experiments (Fig. 2b, 2d).

Not only artificial manipulation of *R* gene expression and dose but also zygosity can affect the mode of immunity. HR induction was also observed for the *Tm-2* gene of tomato and its allelic gene, *Tm-2^2^*, conferring ER to tomato mosaic virus (ToMV). The *Tm-2* and *Tm-2^2^* confer extreme resistance with little or no induction of HR to ToMV in a homozygous state but induce moderate resistance with accompanying HR-like cell death in a heterozygous state or even in a homozygous state at high temperatures (Hall 1980) suggesting that *Tm-2* and *Tm-2^2^* are involved in cell death induction.

We also showed that *Ry_sto_*’s mode of action, unusually for a TIR-NLR, is temperature independent (Fig. 4b, c). Many *R* genes are non-functional at 28°C or above. This was described for the tobacco *N* gene, the *Mi-1* gene in tomato conferring resistance to root-knot nematodes (Jablonska et al., 2007), the *Ny-1* gene conferring resistance to PVY (Szajko et al., 2014), the NLR gene *SNC1* in Arabidopsis (Yang and Hua 2004) and many others. All these genes induce defense in permissive temperature but when shifted to elevated temperature the HR is arrested. In contrast, we have shown that *Ry_sto_-* mediated immunity is still effective in different *Solanaceae* species at or above 28°C (Fig. 4b, 4c). The exact mechanism of temperature sensitivity remains elusive. It might be dependent on the transcription factor PIF4 which acts as a negative regulator of plant immunity and modulation of its function, which alters the balance between growth and defense (Gangappa et al., 2016). It may also relay on Enhanced Disease Susceptibility1 (EDS1) and Phytoalexin Deficient 4 (PAD4)-transcript accumulation which together promote basal resistance and plant growth inhibition in immune-activated backgrounds (Rietz et al., 2011).

A detailed understanding of the functional relationship between HR and ER has not been yet proposed. In principle, the genes for ER act earlier and more efficiently than the genes for HR, which was shown in comparative studies on the potato *Rx* and *N* genes (Bendahmane et al. 1999). *Ry_sto_-*mediated ER is epistatic to *Ny-1* mediated HR against PVY in potato (Fig. 4). We also provide evidence that *Ry_sto_/Ny-1* plants are PVY free at elevated temperature (28°C) in contrast to *Ny-1* plants (Baebler et al., 2014). Collectively, *Ry_sto_*-mediated ER is epistatic to HR. Nevertheless, there is a possibility that ER and HR pathways might be interdependent. For instance, Ry_sto_ might have been competitive with Ny-1 in recruitment of some signaling components. As Ry_sto_ may have higher affinity to selected components than Ny-1, *Ry_sto_*- mediated immunity is stronger and goes quicker. These components might act as helper NLRs especially since R proteins also engage in complex genetic and biochemical interactions (Kamoun et al., 2018).

To date 314 *R* genes have been characterized (Kourelis et al., 2018). Only 128 have a proposed mode of action, based on an even lower number of detailed studies of molecular mechanisms. The underlying mechanisms of the remaining 186 *R* genes are yet unknown (Kourelis et al., 2018). In the *Solanaceae* CNLs outnumber TNLs by 4.7: 1 (Witek et al., 2016). Tobacoo *N* mediates recognition of the helicase domain p50 via interaction with NRIP1 protein and activates a resistance response requiring several known general cofactors of disease resistance: EDS1, SGT1, RAR1, HSP90, members of the COP9 signalosome (Liu et al. 2002, 2003, 2004) and NRG1, the CNL helper protein (Peart et al., 2005). Since *Ry_sto_* contains a TIR domain we decided to search for a mode of action similar to N.

In *EDS1* and *NRG1* mutant *N. benthamiana* plants, the HR after PVY infection and *Ry* transient expression is lost compared to HR observed in non-transformed control plants. This clearly shows that *Ry_sto_*’s mode of action is *EDS1-* and *NRG1-* dependent (Fig. 5), while *Pvr-4* mediated ER to PVY infection in *N. benthamiana* is EDS1-independent (Kim et al. 2018).

In our investigation, HR occurred after PVY infection and *Ry_sto_* transient expression in SA-depleted *NahG* plants (Fig. 5). This indicates that increase of SA is not crucial for *Ry_sto_*- mediated immunity (Fig 5). Interestingly *EDS1* is considered to be a positive SA regulator (Gangappa et al. 2016). Moreover, the EDS1/PAD4-controlled regulatory feedback loop involving SA contributes to the suppression of plant immune response at elevated temperature (Wang et al., 2009). This is in line with a reported lowering of EDS1 protein levels under high temperature conditions (Stuttman et al., 2009). As resistance after PVY infection in *Ry_sto_* plants was still observed both in 28°C and 32°C we concluded that neither *EDS1* nor *NRG1* are involved in temperature sensitivity. Moreover, *EDS1* may also be involved in a signaling pathway unrelated to SA.

Not only *NRG1* but also *ADR1* activity may contribute to TNL-mediated immunity (Dong et al. 2016), so additional studies on *ADR1* involvement in *Ry_sto_*’s mode of action are required.

We propose *Ry_sto_*-mediated immunity to ER against PVY results from *Ry_sto_* recognition of PVY coat protein and recruitment of such downstream signaling components as *NRG1* and *EDS1.* Conceivably, components of the co-chaperone complex: SGT1, RAR1, HSP90 are also involved in *Ry_sto_* mediated immunity as factors supporting correct folding of resistance proteins (Peart et al. 2005). *Ry_sto_* might also be negatively regulated through miRNA activity either by promoting mRNA degradation, inhibiting translation, or suppressing transcription by epigenetic modification.

In summary, our results demonstrate that *Ry_sto_* plays an important role in defense, and may prove valuable for breeding PVY-resistant cultivars of potato and other *Solanaceae* crops. Why some NLRs trigger ER and some HR remains obscure, but nevertheless, this type of recognition leads to durable and efficient resistance.

## MATERIAL AND METHODS

### Plant material

All potato cultivars (*Solanum tuberosum* ssp. tuberosum) and viruses: PVY^0^, PVY^N^, PVY^N-Wi^ and PVX, PVA were obtained from Laboratory for Potato Gene Resources and In Vitro Cultures at the Institute of Plant Breeding and Acclimatisation - National Research Institute, Bonin (Poland). Plants were grown for four weeks in soil under controlled environmental conditions (22°C; 16 h light, and 18°C 8 h dark) as described previously (Szajko et al. 2008). Tobacco plants *Nicotiana tabacum* cv. Xanthi-nc and *Nicotiana benthamiana* were grown for six weeks in soil under controlled environmental (22°C, 16h light, 8h dark) as described previously (Hoser et al. 2013). Transgenic potato and tobacco plants were regenerated followed Agrobacterium leaf disc transformation as described by Mac et al. (2004).

### Mapping population

Diploid potato (2*n* = 2*x* = 24) mapping population was developed by a cross between PVY-resistant heterozygous dihaploid clone dH Alicja, and susceptible clone 83 −3121. dH Alicja was obtained from potato cultivar Alicja via parthenogenesis. The ER to PVY in dH Alicja was conferred by the gene *Ry_sto_* which was derived from clone *MPI 55.957/54*. *MPI 55.957/54* had in its pedigree *Solanum stoloniferum* from wild *Solanum spp*. collection of Max Planck Institute (Ross H. (1958) *Handbuch der Pflanzenzüchtung*, 2^nd^, Vol. III:106- 125).

### Short- and long-read RenSeq and bioinformatic analysis of the sequencing data

Short-read RenSeq libraries and enrichment was carried out on susceptible (S, 83-3121 clone), resistant (R, dhAlicja) and 149 bulked susceptible plants (BS) as described (Jupe,. 2013 and Witek et al., 2016). Enriched libraries were sequenced with Illumina MiSeq platform using 500 cycles. Additionally, RenSeq was performed on cDNA from resistant parent as described previously (Witek et al., 2016) and sequenced on Illumina HiSeq platform with 300 cycles. Long-read PacBio RenSeq was performed as described in Witek et al. (2016) and PacBio was sequenced on a single RSII SMRT cells what resulted in 69,737 Reads of Interest (ROI) with minimum three passes and accuracy above 90%. and are available under accession number PRJEB28862. Reads obtained from SMRT RenSeq were assembled using Geneious 8.1.2 software as described before (Witek et al.,2016) resulting in 1,555 contigs. R, S and BS reads from Illumina 250PE were mapped to contigs derived from SMRT RenSeq assembly using BWA (Li and Durbin, 2009), with default settings. SNP calling and candidate prediction was performed as described previously (Jupe et al.,2013, Witek et al.,2016). Contigs showing >95% of susceptible allele in mapped BS data were considered as linked. Additionally, candidate NLRs showing presence/absence polymorphism between R and S parent, and linkage to resistance based on BS samples were called. Briefly, numbers of pair-end mapped reads to contigs derived from SMRT RenSeq assembly were calculated using TSL Galaxy built-in scripts (MacLean and Kamoun, 2012) for each R, S and BS samples. Resulting data were sorted and visualized in Microsoft Excel software. As a presence/absence polymorphism, contigs with at least 250 mapped reads from R parent, and less than 10% and 18% of that number for S and BS samples, respectively, were considered. Expression of the candidate genes was determined using cDNA-RenSeq data from R parent as described previously (Andolfo et al., 2014, Witek et al., 2016).

### DNA extraction

RenSeq experiments and PCR reactions were conducted on gDNA freshly extracted from young leaves (both MiSeq and PacBio protocols) using the DNeasy Plant Mini Kit (Qiagen, Hilden, Germany) according to the manufacturer’s protocol.

### RNA extraction for RenSeq experiments

For the cDNA RenSeq experiment, RNA was extracted using TRI-Reagent (Sigma-Aldrich, MO, USA) and Direct-zol RNA MiniPrep Kit (Zymo Research, CA, USA), following manufacturer’s recommendations. First-strand cDNA was made using a mix of oligo-dT and random hexamer primers and SuperScript II First-Strand Synthesis System (Sigma-Aldrich, MO, USA). The second strand was made as described (Rallapalli et al., 2014).

### Cloning of the candidate ORFs

For each of 12 selected PacBio contigs PCR primers flanking predicted ORFs were designed. All primers were supplemented with specific 5′ and 3′ extensions to make them compatible with custom USER expression vectors used in this study as described previously (Witek et al., 2016) (see Supplementary Primers Table2). Candidate genes were PCR-amplified from R parent gDNA in 25-μl PCR reactions (35 cycles with annealing at 62 °C and 7 min extension at 72 °C) using Kappa HiFI HotStart Uracil+ Fidelity Polymerase (Manufacturing, R&D Cape Town, South Africa). 30 ng of purified PCR product was hybridized with 30 ng pICSLUS0003 vector in the presence of 1 μl of USER enzyme mix (New England Biolabs, Inc., MA, USA) as described (Witek et al., 2016). All constructs were verified by DNA sequencing. Plasmids containing the candidate genes were transformed into *Agrobacterium* strain GV3101 for transient complementation assays or LBA404 for a stable potato using the method described (Mac et al.,2004). To create a genomic construct of *Ry_sto_* (*c630*) under its native regulatory elements, we PCR-amplified the whole contig (7.5 kB) that was assembled from PacBio reads. PCR amplicons were hybridized into USER - vector pICSLUS0001 lacking 35S promoter and OCS terminator.

### Transient expression assay

Infiltrations of *Agrobacterium* GV3101 carrying the plasmids containing candidate genes were performed in *N. benthamiana*. Bacteria were grown in YEB medium (5 g of beef extract, 5 g of bacteriological peptone, 5 g of sucrose, 1 g of yeast extract, and 2 ml of 1 M MgSO_4_ in 1 litre of milli-Q water) supplemented with gentamicin, rifampicin and kanamycin for all constructs. After 1 day, overnight cells were centrifuged at 3,500 rpm and resuspended in infiltration buffer containing 10 mM MgCl_2_, 10 mM MES (pH 6.5), 100 μM acetosyringone to a final OD600 of 1 and infiltrated into leaves of 4-week-old plants with a 3-ml syringe. HR reaction was observed between 2 and 3 day after infiltration. The same method was used for bacteria carrying a binary plasmid pBIN53 with PVY sequences. Each experiment was repeated and included at least three independent biological replicates.

### Resistance assay

In all experiments related to monitoring of PVY infection the isolate NIB-NTN (GenBank: AJ585342.1) was used. The infection was performed in each time on a seven-week-old *N. tabacum* plants or 4-week-old potato plants under conditions as described (Baebler et al. 2014). In experiments related to different PVY isolates plants were infected under the same experimental conditions with following PVY isolates: 0 (AJ890349), N (FJ666337), N-Wilga (EF558545), or unrelated viruses PVA, PVX and TMV(U1).

### RNA extraction and gene expression analysis (qPCR)

Total RNA was extracted using a modified guanidinium-phenol-chloroform protocol (Chomczynski and Sacchi, 1987), treated with DNase1 (ThermoFisher Scientific), and subjected to reverse transcription using mix of random hexamer and oligo primer and RevertAid First-Strand cDNA Synthesis Kit (ThermoFisher Scientific). Gene expression analysis via RT-qPCR was performed using the LightCycler^®^480 instrument and LightCycler^®^480 SYBR Green I Master Kit (Roche). Relative gene expression levels were determined using a standard curve method and the value for each target gene was then normalized against the mean of expression values of two reference genes: *EF1* and *L23* or *EF1* and *Sec3* for *N tabacum* and *N. benthamiana* or potato cultivars respectively, as described (Liu et al., 2012, Tang et al. 2017). Measurements for each sample was analysed in four technical replicates and two dilutions. All primers used for qPCR are described in Supplementary Table 4.

### PVY proteins cloning

A full-length infectious clone PVY-N605 (Jakub et al. 1997) was used to create all PVY constructs. Each putative PVY protein was PCR amplified and cloned to pBIN53 vector which carries the CaMV 35S promoter, the PVY 5′ and 3′ UTRs separated by a polylinker, and a poly(A) tail vector as described previously (Mestre et al., 2000).

### Western blotting

Agro-infiltrated leaves were collected and ground in liquid nitrogen. Total protein was extracted by incubating the ground leaf samples in extraction buffer containing 100 mM Tris HCl pH 8.0, 1 mM EDTA, 150 mM NaCl, 7.2 mM β-mercaptoethanol, 0.5 mM 4-(2- aminoethyl) benzenesulfonyl fluoride hydrochloride (AEBSF) and 0.03 l M PMSF (phenylmethylsulfonyl fluoride) for 15 min. The homogenate was centrifuged at 21,000×g for 10 min and the supernatant was collected. Samples were fractionated by 12.5% SDS–PAGE and subjected to immunoblot analysis using appropriate alkaline phosphatase-conjugated anti PVA antibodies (Bioreba, Switzerland). Immunoblots were developed using the NBT/BCIP colorimetric detection kit from BioShop Canada Inc.

### Statistical analysis

Statistical analyses were conducted using R 3.2.2 within R Studio 0.99.483 Technical replicates consist of readings from the same plant in the same experiment, whereas biological replicates consist of independent plants. Data were analysed using the following pipeline: data were assessed for their suitability to be analysed using parametric tests by testing for the normal distribution of the residuals and Shapiro–Wilk test. If the data were suitable for conducting parametric tests, then analysis of variance (ANOVA) were used. As post-hoc the Dunnett’s or Tuckey’s HSD test was used.

## ACKNOWLEDGMENTS

We would like to thank our co-workers, in particular Dr. Magdalena Krzymowska, for critical comments on the manuscript, Prof Ewa Zimnoch-Guzowska for discussions and help in the preliminary phase of the project. We would like to acknowledge Danuta Strzelczyk-Żyta and Izabella Barymow for the technical support, Baptiste Castel for sharing seeds of *nrg1 N. benthamiana* knockout plants and Brian Staskawicz for sharing seeds of *eds1 N. benthamiana* knockout plants.

**Author contributions:**
M.G-B., K.W and J.H. performed most of the experimental work, data analyses, and writing; A. W., I.W-F., H.J., K.S. and K.M performed the research; J.J. and W.M. edited the article; J.H. supervised the work and assisted with writing.

